# Epidermal Resident Memory T Cell Fitness Requires Antigen Encounter in the Skin

**DOI:** 10.1101/2025.03.31.646438

**Authors:** Eric S. Weiss, Toshiro Hirai, Haiyue Li, Andrew Liu, Shannon Baker, Ian Magill, Jacob Gillis, Youran R. Zhang, Torben Ramcke, Kazuo Kurihara, The ImmGen Consortium OpenSource T cell Project, David Masopust, Niroshana Anandasabapathy, Harinder Singh, David Zemmour, Laura K. Mackay, Daniel H. Kaplan

## Abstract

CD8^+^ tissue resident memory T cells (T_RM_) develop from effectors that seed peripheral tissues where they persist providing defense against subsequent challenges. T_RM_ persistence requires autocrine TGFβ transactivated by integrins expressed on keratinocytes. T_RM_ precursors that encounter antigen in the epidermis during development outcompete bystander T_RM_ for TGFβ resulting in enhanced persistence. ScRNA-seq analysis of epidermal T_RM_ revealed that local antigen experience in the skin resulted in an enhanced differentiation signature in comparison with bystanders. Upon recall, T_RM_ displayed greater proliferation dictated by affinity of antigen experienced during epidermal development. Finally, local antigen experienced T_RM_ differentially expressed TGFβRIII, which increases avidity of the TGFβRI/II receptor complex for TGFβ. Selective ablation of *Tgfbr3* reduced local antigen experienced T_RM_ capacity to persist, rendering them phenotypically like bystander T_RM_. Thus, antigen driven TCR signaling in the epidermis during T_RM_ differentiation results in a lower TGFβ requirement for persistence and increased proliferative capacity that together enhance epidermal T_RM_ fitness.

## Introduction

Tissue resident CD8 memory T cells (T_RM_) are a highly abundant, non-circulating, long-lived subset of memory T cells that play an important role in protecting against re-infections [1, 2]. T_RM_ also provide immunosurveillance against neoplasia and are thought to be pathogenic in some autoimmune diseases [3–9]. In the skin, following infection with vaccinia virus (VV) or herpes simplex virus, T_RM_ develop from CD8^+^ T cell effectors (T_EFF_) that expand following priming in the lymph node and are recruited into the skin by inflammatory signals where they preferentially reside in the epidermis [1, 10–15]. Unlike T_RM_ in some tissues, re-encounter with cognate antigen in the skin is not required for T_RM_ differentiation [16–19]. Epidermal T_RM_ that encounter cognate antigen in the skin or bystander T_RM_ that do not encounter antigen both develop with comparable efficiency and both express similar levels of the canonical T_RM_ markers CD103 and CD69[20, 21].

The cytokine transforming growth factor β (TGFβ) is required for cutaneous T_RM_ development at multiple stages. TGFβ signaling in the lymph node during steady state epigenetically preconditions naïve CD8 T cells allowing for later T_RM_ differentiation [22]. T_EFF_ recruited into skin require TGFβ signaling for entry into the epidermis and differentiation into T_RM_ [23, 24]. Once epidermal T_RM_ have differentiated, these cells continue to require TGFβ signaling to retain epidermal residence. TGFβ is produced bound to the latency associate peptide that prevents bioactivity until the complex is activated which in the epidermis is mediated exclusively by activation via the integrins α_v_β_6_ and α_v_β_8_ expressed by keratinocytes [25–28]. Epidermal persistence of T_RM_ depends on autocrine TGFβ which is transactivated by the integrins ⍺vβ6 and ⍺vβ8[21, 29]. Inducible ablation in T_RM_ of a required component of the TGFβ receptor (TGFβRII) or TGFβ1 as well as ablation of ⍺vβ6 and ⍺vβ8 in keratinocytes all result in loss of epidermal T_RM_ [21, 30].

We previously reported that local antigen experienced T_RM_ that have encountered cognate antigen in the skin are better able to persist in the epidermis than bystander T_RM_ that have not encountered antigen in the skin when active TGFβ is limited, despite the fact that both groups of cells had prior activation within the lymph node [21]. This is evident when TGFβ activation is experimentally reduced by small molecule inhibition of ⍺vβ6 and ⍺vβ8 and when established T_RM_ compete with newly recruited T_EFF_ cells for limiting amounts of TGFβ activation. In both instances, bystander T_RM_ are preferentially lost from the epidermis while local antigen experienced T_RM_ persist. Thus, encounter with antigen in the skin results in more fit T_RM_ and represents a potential opportunity to preferentially enrich antigen-specific over bystander T_RM_ cells during repeated challenges.

Herein, we delineate mechanisms of T_RM_ homeostasis by demonstrating that within a population of endogenous T_RM_, antigen encounter in the skin is required for their full differentiation whereas bystander T_RM_ are maintained at an earlier developmental stage. We also find that local antigen experienced T_RM_ have increased proliferative capacity during a recall response compared to bystander T_RM_ that is dependent on affinity of antigen experienced in skin thus providing an additional functional attribute defining T_RM_ fitness. Finally, we show that expression of TGFβRIII by local antigen experienced T_RM_ which can be induced following TCR ligation is required for epidermal persistence when active TGFβ is limiting.

## Results

### Epidermal T_RM_ are transcriptionally heterogeneous

The recruitment of effector CD8^+^ T cells into mouse flank skin via a viral infection (e.g. Vaccinia Virus) or through an inflammatory stimulus (e.g. “DNFB-pull”) results in comparable numbers of long-lived CD103^+^ epidermal T_RM_ and is independent of the presence or absence of cognate antigen in the skin [18, 29, 31]. This, however, has only been demonstrated using HSV-specific (gBT-I) or OVA-specific (OT-I) TCR transgenic T cells. To determine if endogenous polyclonal CD8^+^ T cells share this phenotype, we employed a dual VV infection “DNFB-pull” model. Cohorts of wildtype C57BL/6 mice were infected on the left flank with VV by skin scarification (**Figure 1A**). The infection expanded CD8^+^ effectors specific for VV antigens which were then recruited into the left flank in response to the ongoing infection-induced inflammation. DNFB is applied to the right flank 5 days post infection to recruit into the skin circulating CD8 effectors expanded by infection. Mice were then rested for 50+ days to allow for the formation of T_RM_. We have previously found that evaluation of T_RM_ numbers in the epidermis by flow cytometry is much less accurate than direct immunofluorescent visualization [21]. Evaluation of whole mounted epidermal sheets stained with anti-CD8 revealed comparable numbers of CD8^+^ T cells at the VV infected (left flank) and DNFB treated (right flank) sites (**Figure 1B and C**). Expression of CD103 on T_RM_ as evaluated by flow cytometry was also similar at both sites (**Figure 1D and S1A**). To evaluate the frequency of antigen-specific T cells at each site, skin from a separate cohort was examined by flow cytometry using the B8R tetramer which recognizes an immunodominant epitope of VV in H2-K^b^[32]. We observed equivalent numbers of B8R^+^ T_RM_ at the VV-infected and DNFB treated sites (**Figure 1E and F**). From these data we conclude that polyclonal CD8 T cells form epidermal T_RM_ with comparable efficiency when recruited into the skin by viral infection or sterile inflammation and that the presence or absence of cognate antigen in the skin has no effect on the number or frequency of antigen-specific T_RM_.

**Figure 1.**
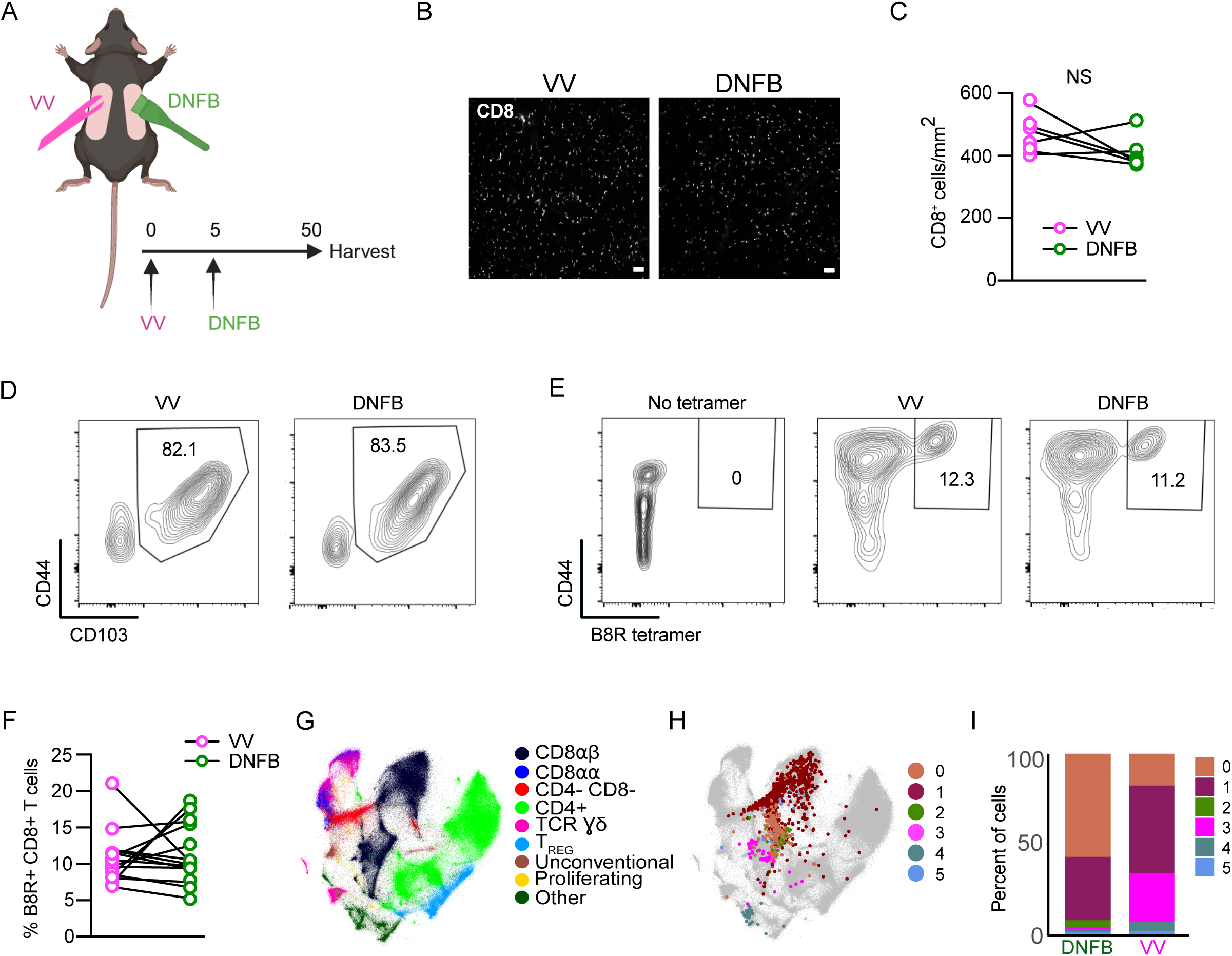
Epidermal T_RM_ are transcriptionally heterogeneous. (A) Experimental design. Mice were treated with vaccinia virus infection by skin scarification on the left flank on day 0, and then on day 5 post infection the right flank was painted with 0.15% DNFB. On day 56, flanks were harvested for either epidermal whole mount, flow cytometry, or cells were sorted for scRNA-seq. (B) Representative images and (C) quantification of epidermal whole mounts of VV and DNFB treated flanks harvested on day 50 post infection stained for CD8a. (D) Representative flow plots gated on CD45^+^ CD3^+^ CD8^+^ CD90.2^+^ cells isolated from VV or DNFB treated flanks. (E) Representative flow plots and (F) quantification of B8R tetramer binding of CD103+ T_RM_ gated as in (D) isolated from VV or DNFB treated flanks. (G) The integrated Minimal-Distorted Embedding of all 96 experiments from the ImmgenT consortium with annotated clusters. (H) Minimal-Distorted embedding visualization of transcriptional clusters of skin T_RM_ projected over the ImmgenT dataset. (I) The percentage of cells in each transcriptional cluster found in VV or DNFB treated sites. Each symbol represents paired data from the same individual animal (C, F). Data shown to be nonsignificant by paired Student’s t-tests (C,F). Data are representative of 3 separate experiments. Scale bar in (B) represents 50um.

Although T_RM_ efficiently populate the epidermis independently of cutaneous cognate antigen, we have previously demonstrated that T_RM_ that form at skin sites containing cognate antigen are functionally distinct from bystander T_RM_ that form in its absence[21]. To analyze potential transcriptional differences between local antigen experienced and bystander T_RM_, we performed single-cell RNAseq in collaboration with the ImmgenT consortium [33]. The consortium consists of multiple groups that isolated populations of murine T cells from multiple tissues and contexts and then subjected them to single-cell RNA-seq [33](see Methods). 627,692 mature T cells from 80 different experiments and 703 samples were integrated and batch-corrected using totalVI [34] (see Methods).

Cells were projected in two dimensions using Minimal-Distorted Embedding (see Methods) revealing the expected transcriptional clustering of distinct types of T cells (**Figure 1G**). As part of the consortium, we isolated cells from the flanks of rested (>65 days) VV-infected or DNFB treated flank skin by enzymatic digestion. Cells were pooled from 10 mice and purified based on expression of CD90 and CD8 (**FigureS1B**). As expected, most cells expressed CD69 and CD103 (**Figure S1C**). Skin cells from each group and naïve spleen cells were hash tagged and analyzed by single-cell RNA-seq.

Analysis of transcripts revealed a total of 6 clusters (0-5) which are shown overlayed on the complete ImmgenT cell embedding (**Figure 1H, S1D**). As expected, most cells clustered with CD8⍺β T cells (**Figure 1G, black**). There was, however, a high degree of heterogeneity and some, possibly contaminating cells fell outside of this region. A comparison of the relative frequency of cells in each of the clusters (0-5) isolated from the VV or DNFB sites revealed a strong enrichment of cluster 3 in the VV group and of clusters 0 and 2 in the DNFB group (**Figure 1I, magenta**). This suggested that cluster 3 cells were manifesting a distinct transcriptional state likely induced by antigen encounter in the skin.

### Cutaneous antigen is required for complete T_RM_ differentiation

To test the hypothesis that cells in cluster 3 could be T_RM_ that have encountered antigen in the skin during development, we performed signature score enrichment analysis comparing the transcriptional profile of our cells to published core genes of well-defined CD8 T cell states. Signature scores were calculated based on normalized differential gene expression compared to core genes of T_RM_ generated by acute infection models (GSE47045), activated T cells (T_ACT_, GSE10239), circulating memory T cells (T_MEM_, GSE41867), and exhausted T cells (T_EX_, GS41867) [35–38]. Notably, cells in cluster 3 showed enriched expression of T_RM_ core genes compared to all other clusters (**Figure 2A**). We performed a similar analysis using pseudo-bulk analysis of all cells isolated from the VV or DNFB sites. Consistent with the enrichment of cluster 3 in cells from the VV site, we found a strong enrichment of T_RM_ core genes in the VV group (**Figure 2B**). These data suggest that cells in cluster 3 isolated from the VV site represent fully differentiated T_RM_.

**Figure 2:**
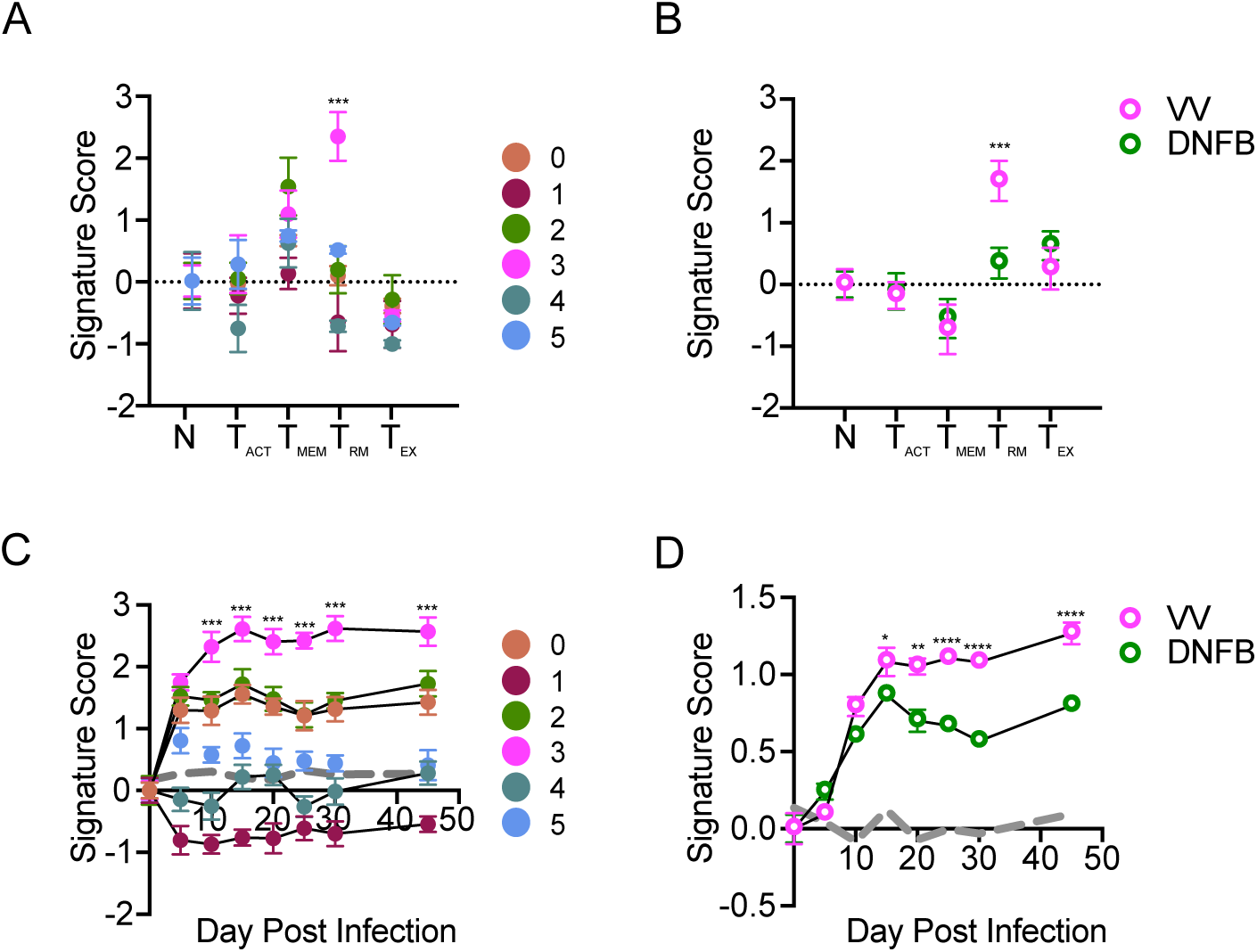
Cutaneous antigen is required for complete T_RM_ differentiation. (A) Signature score analysis of skin T cell clusters calculating the enrichment for previously published T cell state gene sets (N=naïve). Signature score = Mean (average upgenes z-scores) - Mean (average downgenes z-scores). *** p < 0.001 by Dunnett’s multiple comparisons of cluster 3 to all other clusters. (B) Pseudobulk analysis comparing signature scores of cells isolated from VV and DNFB sites as in (A). ** p < 0.01 and *** p < 0.001 by paired Student’s t-tests. (C) Signature score analysis of individual clusters or (D) cells isolated from VV and DNFB sites calculating the enrichment for compared to epidermal T_RM_ isolated at the indicated day post infection. ***p < 0.001 by Dunnett’s multiple comparison test of cluster 3 to all other clusters in (C). * p < 0.05, ** p < 0.01, **** p < 0.0001 by paired Student’s t-tests in (D). Gray line represents a randomized control generated by the average enrichment of each group compared to a randomly generated gene set of an equal number of probes. Datapoints and error bars represent mean and 95% confidence interval of the relative enrichment of each set of DEGs.

To further test the hypothesis that cluster 3 represents fully differentiated T_RM_, we performed signature score enrichment analysis of clusters 0-5 against a dataset of skin T cells isolated at different times following skin VV infection (GSE79805) [39]. Cells from clusters 0, 2 and 3 showed similar signature scores in comparison with skin T cells up to day 5 post infection, however clusters 0 and 2 then plateaued suggesting a lack of continued differentiation (**Figure 2C**). In contrast, cluster 3 transcripts scored higher at later time points in T_RM_ differentiation. Signature scores for cells isolated from the DNFB and VV sites were similar at early time points but diverged at later time points with increased scores observed in cells from the VV site (**Figure 2D**). Taken together, these data suggest that T_RM_ isolated from skin sites where they can encounter cognate antigen are transcriptionally more fully differentiated T_RM_ while bystander T_RM_ isolated from a site lacking cognate antigen remain in a less differentiated state.

### Local antigen experienced T_RM_ have improved expansion in a recall response

Analysis of the 10 highest differentially expressed genes (DEGs) in cluster 3 (**Figure S1D**) showed increased expression of genes associated with T cell activation including *Dusp1* and *Nr4a1* as well as the AP-1 family members J*unb* and *Fos* [40–44]. Given the importance of AP-1 family members in T cell proliferation and differentiation, we next tested whether this corresponded to functional differences in T_RM_ activity [43, 45–48].

An important functional property of T_RM_ is their capacity to rapidly expand following re-encounter with cognate antigen [49, 50]. To determine whether local antigen experienced and bystander T_RM_ have different recall responses, we repeated our dual VV infection and DNFB “pull” model in combination with an OT-I adoptive transfer model to allow for antigen recall using the SIINFEKL peptide. CD90.1^+^ OT-I cells were adoptively transferred into naïve C57BL/6 mice followed by infection with a vaccinia virus expressing SIINFEKL peptide, OVA_257-264_ (VV-OVA) on the left flank. On day 5 post infection, the right flank was treated with DNFB to recruit expanded OT-I effectors to the site (**Figure 3A**). On day 50+ after infection, topical SIINFEKL peptide or control PBS was painted on both flanks in a single application (1° recall). The total number of OT-I cells in the epidermis 6 days later was determined by immunofluorescent microscopic evaluation of the CD90.1 congenic marker in epidermal whole mounts (**Figure 3B-C**). Restimulation with peptide led to an expansion of T_RM_ at both sites compared to PBS treated controls. Notably, local antigen experienced OT-I T_RM_ cells in the VV-OVA treated flanks expanded to a larger extent compared to bystander T_RM_ in DNFB treated flanks. To inhibit potential recruitment of circulating effector OT-I cells, we repeated these experiments administering FTY720 to block migration of T cells from lymph node or titrated anti-Thy1.1 antibody to ablate circulating OT-I but not T_RM_ (**Figure 3D**). Both methods successfully depleted OT-I from the blood (**Figure S2A-D**), without affecting T_RM_ in the skin (**Figure S2E**). Administration of FTY720 (**Figure 3E-F**) or anti-Thy1.1 treatment (**Figure 3G**) had no effect on the number of OT-I T_RM_ in the epidermis after a 1° recall response and local antigen experienced T_RM_ from the VV-OVA treated flank still expanded to a greater degree than bystander T_RM_ from the DNFB treated flank. Thus, the increased number of epidermal OT-I at the local antigen experienced VV site during a 1° recall does not appear to result from recruitment of circulating cells.

**Figure 3.**
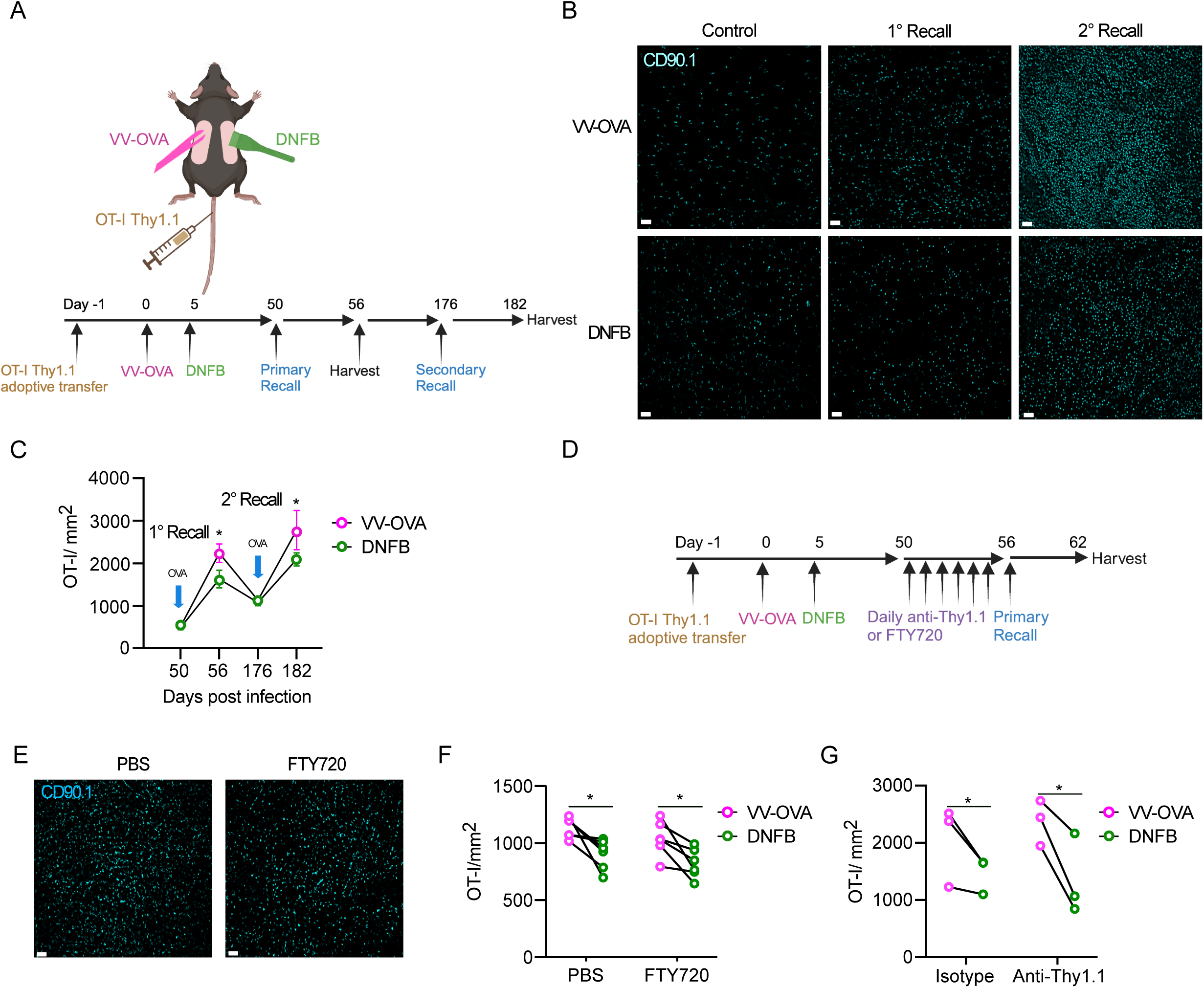
Local antigen experienced T_RM_ have improved expansion in a recall response. (A) Experimental design. Mice were adoptively transferred with Thy1.1^+^ OT-I T cells on day −1, then infected with OVA-expressing vaccinia virus by scarification on the left flank on day 0. On day 5 post-infection the right flank was painted with 0.15% DNFB. On day 50, a primary recall response with SIINFEKL peptide in acetone and olive oil was painted on both flanks and harvested on day 56. In some cohorts, mice were allowed to rest for an additional 120 days and then treated with topical SIINFEKL peptide again on day 176 and harvested on day 182. (B) Representative images and (C) quantification of epidermal whole mounts isolated from VV-OVA and DNFB treated flanks stained for anti-Thy1.1 (n=10 animals). (D) Experimental design. Mice were treated as in (A), but for 6 days prior to SIINFEKL treatment they were given either i.p. FTY720, i.p. PBS, i.v. titrated anti-Thy1.1 or i.v. isotype control. (E) Representative epidermal whole mounts images and (F) quantification of skin from VV-OVA treated flanks on day 6 of a primary recall response treated with either FTY720 or PBS. (G) Quantification of total Thy1.1 OT-I cells in epidermal whole mounts on day 6 after a primary recall response in mice treated with either isotype or anti-Thy1.1 depleting antibody. Each symbol represents paired data from an individual same animal (F, G). Data are representative of 3 independent experiments. *p<0.05 by paired Student’s t-tests. Unpaired Student’s t-tests between FTY720 treated (F) or Anti-Thy1.1 treated (G) flanks and PBS treated flanks shows non-signficance for both VV-OVA and DNFB, Scale bar represents 50um (B, F). Each symbol represents the mean +/- SEM (C).

Finally, to determine whether increased expansion by local antigen experienced T_RM_ persists with multiple rounds of stimulation, we repeated this experiment but allowed mice to rest for 120 days after the 1° recall response. The number of T_RM_ at both the VV-OVA and DNFB sites contracted down to equivalent numbers that were increased compared to PBS treated, control mice (**Figure 3C**). Mice were then given a 2° recall by a second application of peptide at day +176. The number of T_RM_ expanded but T_RM_ at the VV-OVA site increased by a larger amount than T_RM_ at the DNFB site (**Figure 3B-C**). Based on these data, we conclude that T_RM_ encountering antigen in the skin during development expand more efficiently in response to antigen re-encounter and this phenotype persists long-term with subsequent antigen encounters.

*Local antigen experienced T_RM_ exhibit increased proliferation during a recall response* We next hypothesized that the increased expansion of local antigen experienced T_RM_ during a recall response resulted from increased proliferation. To test this, as above, WT mice adoptively transferred with OT-I cells were infected with VV-OVA on the left flank and DNFB on the right flank 5 days later. After at least 50 days of rest, both sides were painted with topical SIINFEKL peptide and mice were administered 2 mg of BrdU i.p. daily (**Figure 4A**). Epidermal sheets for immunofluorescent visualization were harvested 2 days later. We noted a clear increase in the percent and total number of dividing cells as well as the total number of OT-I T cells in the epidermis at the VV-OVA site compared with the DNFB site (**Figure 4B-D**). Similar results were obtained by flow cytometry, with an increase in the percentage of BrdU^+^ in T_RM_ isolated from the VV-OVA site (**Figure 4E-G**). Staining for Annexin V was equivalent in both groups suggesting that an altered rate of apoptosis does not contribute to the observed changes in expansion (**Figure 4H-I**). In sum, local antigen experienced T_RM_ show augmented proliferation during an antigen recall response compared with bystander T_RM_. Moreover, an enhanced proliferative capacity following antigen restimulation can now be added to other parameters of T_RM_ fitness including enhanced epidermal persistence when active TGFβ is limited.

**Figure 4.**
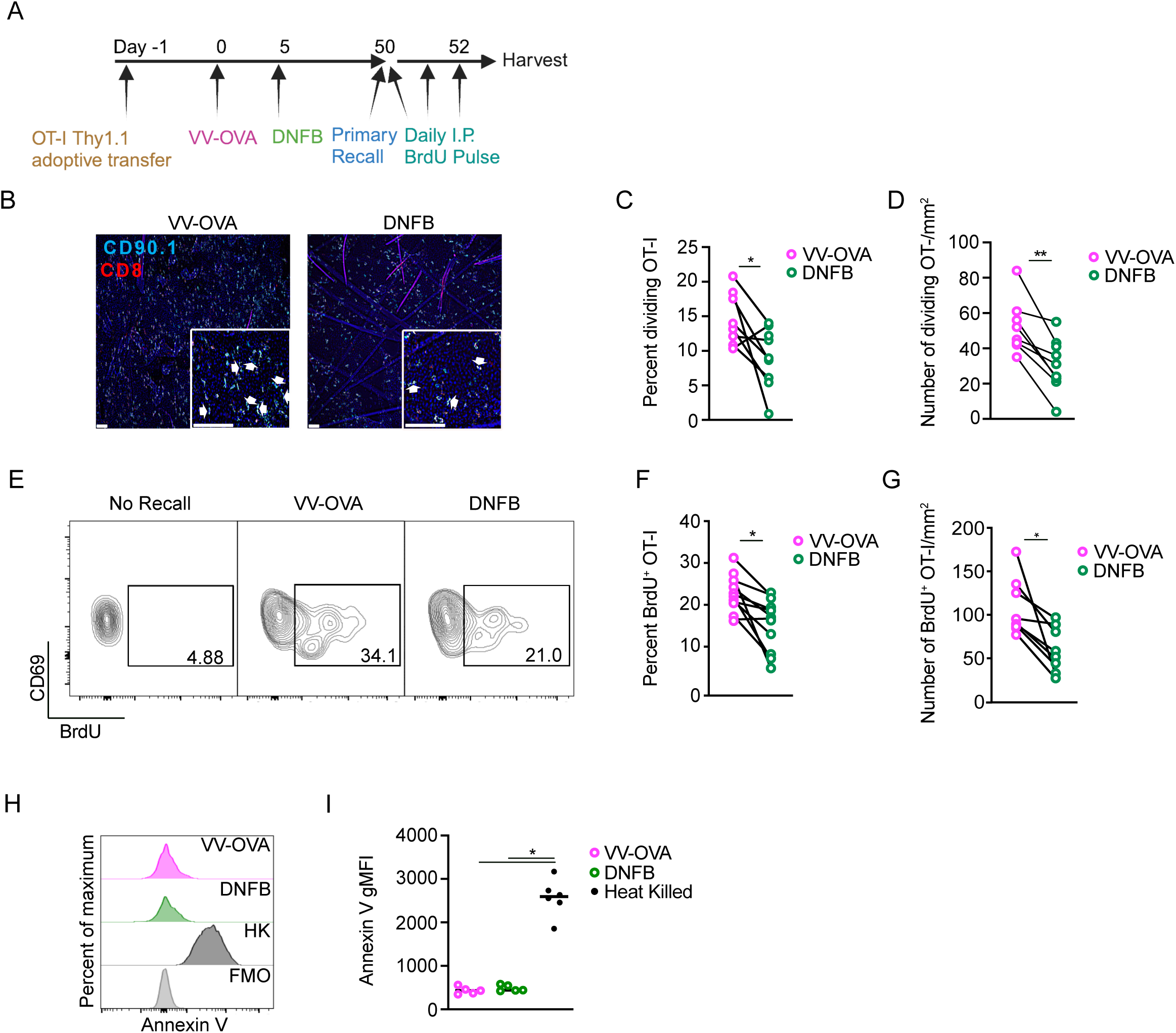
Local antigen experienced T_RM_ have increased proliferation during a recall response. (A) Experimental scheme. (B) Representative images of epidermal whole mounts of VV-OVA or DNFB treated flanks on day 2 of a recall response. Arrows highlight cell doublets. (C) Quantification showing percent OT-I cells that are dividing in epidermal whole mounts on day 2 after primary recall response, and (D) total number of dividing OT-I cells. (E) Representative flow cytometric plots and (F) quantification showing BrdU incorporation in gated CD45^+^ CD8^+^ CD90.1^+^ CD103^+^ CD69^+^ cells isolates from VV-OVA or DNFB treated flanks. (G) Quantification of total numbers BrdU^+^ OT-I cells combining OT-I numbers with percentage of BrdU incorporation in (F). (H) Representative histograms and (I) quantification of Annexin V expression in OT-I cells isolated from VV-OVA or DNFB treated flanks or OT-I cells heat-treated at 60°C for 1 hour (HK). Data are representative of 3 separate experiments. *p<0.05, **p<0.01 by Students paired t-tests (C), (D), (F) and (G) or Dunnett’s test (I). Scale bar represents 50um.

### T_RM_ fitness depends on TCR signal-strength

We next hypothesized that the strength of TCR signal provided in the skin during T_RM_ differentiation should correlate with T_RM_ phenotype at late time points. To generate T_RM_ with a range of local antigen encounter signal strengths, we wanted to expose newly-recruited T cells to either topical SIINFEKL peptide (N4), or a topical altered peptide ligand (APL) SIYNFEKL (Y3) that has ¼ the avidity for the TCR receptor [51]. We hypothesized that encountering an APL would induce an intermediate functional phenotype between full-strength SIINFEKL and no local antigen encounter. To test this hypothesis, WT mice were adoptively transferred with OT-I cells then infected with VV-OVA on the left flank and 5 days later were challenged with DNFB on the right flank (**Figure 5A**).One day later, the DNFB site was painted with PBS, SII**N**FEKL (N4), or an APL SI**Y**NFEKL (Y3), After 50+ days of rest, all mice were challenged in a 1° recall response with topical N4 or control PBS at the DNFB sides (**Figure 5A**). Epidermal sheets for immunofluorescence were harvested 6 days later. As expected, mice restimulated with PBS showed equivalent numbers of T_RM_ regardless of which peptide was provided during development (**Figure 5B-C**). Following a 1° recall response, T_RM_ exposed to N4 (DNFB + N4) expanded to a level equivalent to T_RM_ from the VV-OVA site. In contrast, T_RM_ exposed to Y3 (DNFB + Y3) showed a degree of expansion that was intermediate compared to T_RM_ that did not experience antigen in the skin (DNFB+PBS) and those that experienced SIINFEKL (DNFB + N4). Thus, the strength of TCR engagement determines the fitness of T_RM_ based on their capacity to expand during a 1° recall response.

**Figure 5.**
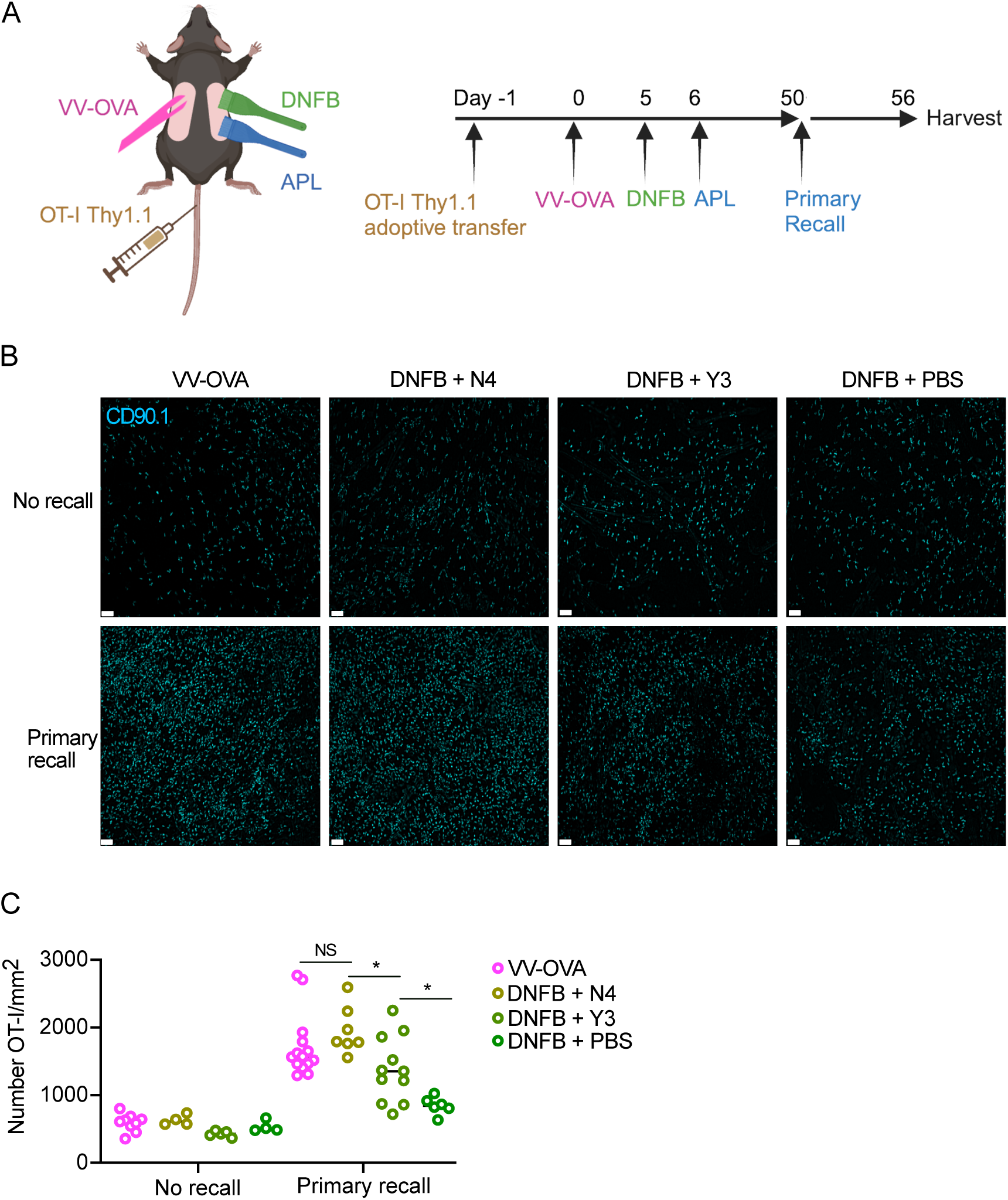
T_RM_ fitness depends on TCR signal-strength. (A) Experimental scheme. Local antigen experienced and bystander T_RM_ were generated and restimulated as previously described, but on the DNFB-treated flanks, altered peptide ligand variants of SIINFEKL were topically applied to the skin 1 day after recruitment to the skin. (B) Representative images and (C) quantification of epidermal whole mounts (CD90.1 cyan) of VV-OVA and DNFB+ APL treated flanks, at steady state (no recall) or 6 days post recall response. Each symbol represents data from an individual animal. Data are representative of 5 independent experiments. *p<0.05 by unpaired Student’s t-tests. Scale bar, 50um.

### Local antigen experienced T_RM_ have improved persistence mediated by TGFßRIII

A feature of local antigen experienced T_RM_ is that they are better able to persist in the epidermis than bystanders when levels of activated TGFβ are limited, either artificially or during competition with newly recruited T_EFF_ [21]. We have previously found that expression of the canonical TGFβ receptors, *Tgfbr1* and *Tgfbr2* have equivalent expression in local antigen experienced and bystander T_RM_ [21]. However, there is a third TGFβ receptor, TGFβRIII, that lacks a signaling component but functions as a reservoir of ligand for TGFβ, increasing avidity of the TGFβ receptor [52, 53]. Notably, expression of TGFβRIII has been reported to increase in T cells following TCR ligation [54]. This was confirmed in vitro with anti-CD3 anti-CD28 stimulated OT-I splenocytes (**Figure 6A**). We also observed increased expression of TGFβRIII in vivo comparing local antigen experienced OT-I T_RM_ with bystanders (**Figure 6B-C**). Based on this data, we hypothesized that increased expression of TGFβRIII on antigen experienced T_RM_ could explain their capacity to maintain epidermal residence when available TGFβ is limiting.

**Figure 6.**
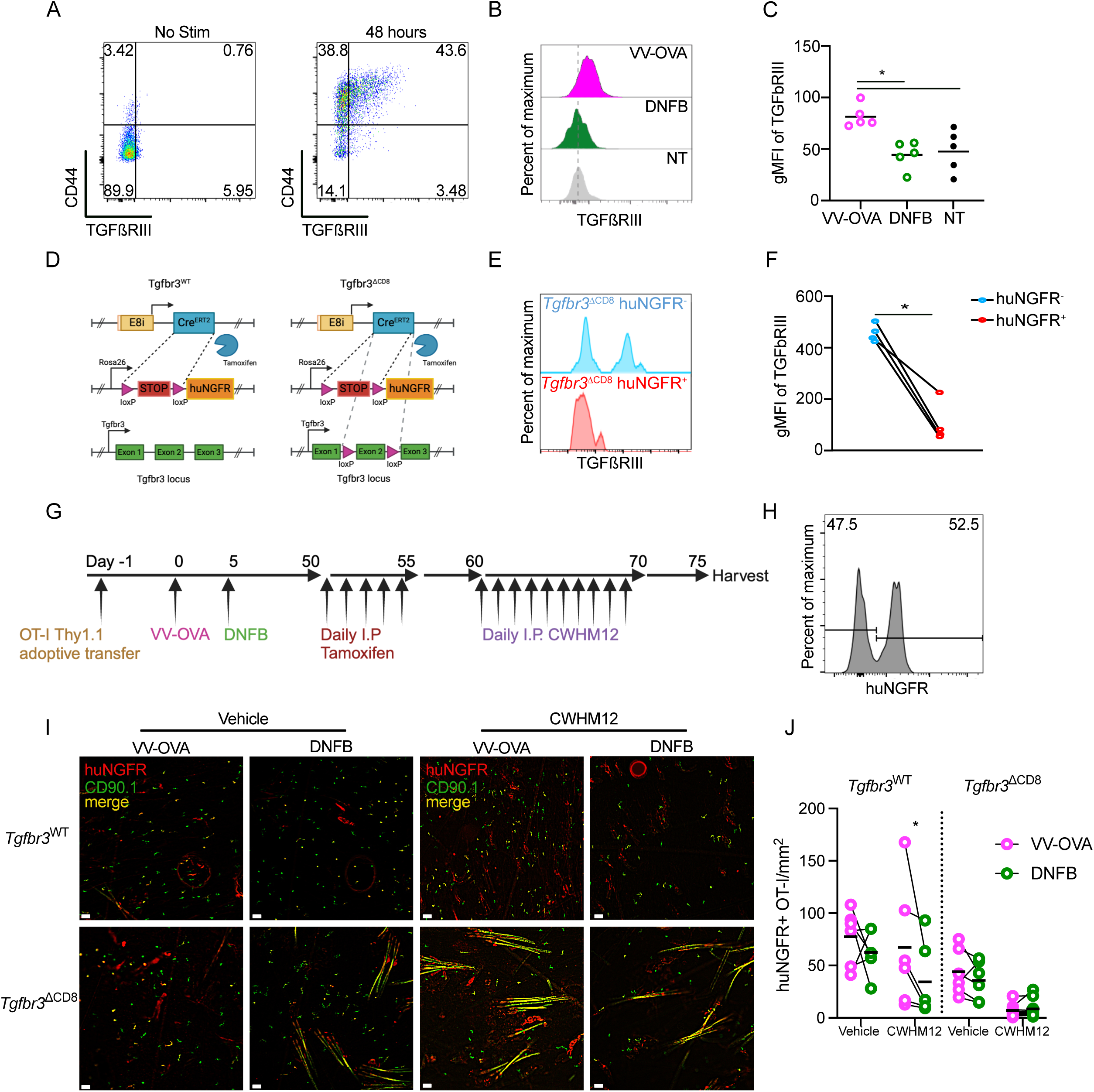
Local antigen experienced T_RM_ have improved persistence mediated by TGFßRIII. (A) Representative flow cytometric plots of TGFßRIII staining of Thy1.1^+^ OT-I cells stimulated in vitro for 48 hours with PBS or anti-CD3 anti-CD28. (B) Representative histograms showing TGFßRIII expressing in CD45^+^ CD3^+^ CD8^+^ CD90.1^+^ gated OT-I cells isolated from VV-OVA, DNFB or untreated flanks at least 50 days post infection. (C) Quantification of (B). (D) Schematic demonstrating genetics of Tgfbr3^WT^ and Tgfbr3^ΔCD8^ mice. (E) Representative histogram and (F) quantification of TGFßRIII expression in huNGFR^-^ or huNGFR^+^ Tgfbr3^ΔCD8^ T cells harvested 5 days following i.p. treatment with tamoxifen then stimulated in vitro for 48 hours with anti-CD3 and anti-CD28. (G) Experimental scheme. (H) Representative histogram of CD45.2+CD3+CD8+CD90.1+ OT-I cells isolated from LNs after tamoxifen treatment demonstrating transformation efficiency. (I) Representative epidermal whole mounts showing Thy1.1 staining (green), huNGFR staining (red) or merge (yellow) of VV-OVA or DNFB treated flanks from mice adoptively transferred with either Tgfbr3^WT^ or Tgfbr3^ΔCD8^ cells, treated with tamoxifen i.p., and then given either PBS or i.p. CWHM12 for 10 days. Hair follicles in the sample present as long yellow streaks. (J) Quantification of total huNGFR^+^ Thy1.1^+^ OT-I in (I). Each symbol represents data from an individual animal. Black lines represent group means. Data is representative of 3 independent experiments. *p<0.05 by Dunnett’s test (C) or paired Student’s t-tests (F) and (J). Unpaired Student’s t-tests between Tgfbr3^WT^ and Tgfbr3^ΔCD8^ vehicle treated groups show non-significance for both VV-OVA and DNFB treated flanks. Scale bar represents 50um.

To test this, we generated OT-I Thy1.1^+^ E8i-cre^ERT2^ huNGFR Tgfbr3^fl/fl^ (Tgfbr3^ΔCD8^) mice which allow for inducible ablation of *Tgfbr3* from T_RM_ as well control OT-I Thy1.1^+^ E8i-cre^ERT2^ huNGFR *Tgfbr3*^WT/WT^ (Tgfbr3^WT^) mice (**Figure 6D**). To validate these mice, Tgfbr3^ΔCD8^ mice were treated with 0.05mg/g tamoxifen i.p. for 5 days. CD8^+^ T cells from tamoxifen treated Tgfbr3^ΔCD8^ mice were then isolated from spleen and lymph nodes and cultured for 2 days with anti-CD3 anti-CD28. Cells were evaluated by flow cytometry for expression TGFβRIII and huNGFR, an indicator of successful induction of cre-mediated excision by tamoxifen. In the huNGFR-negative population where cre was not activated, approximately 50% of the cells expressed TGFβRIII (**Figure 6E-F**). In contrast, TGFβRIII was largely absent from cells co-expressing huNGFR indicative of efficient ablation of *Tgfbr3*.

To ablate TGFβRIII from T_RM_ once they have fully differentiated in the epidermis, we next adoptively transferred CD8^+^ T cells isolated from either Tgfbr3^WT^ or Tgfbr3^ΔCD8^ mice into naïve WT mice followed by skin VV-OVA infection on the left flank. On day 5 post infection, the right flank was treated with DNFB. After at least 50 days of rest to allow for T_RM_ formation, both cohorts were treated daily with 0.05mg/g tamoxifen i.p. to transform approximately 50% of adoptively transferred cells, generating populations of both control huNGFR negative and huNGFR positive OT-I cells. To test the effects of TGFβRIII ablation, TGFβ activation was inhibited for 10 days by i.p. administration of CWHM12, a small molecule inhibitor of integrins ⍺vβ6 and ⍺vβ8 (**Figure 6G-H**).

Analysis of epidermal whole mounts by immunofluorescence revealed that blockade of TGFβ activation reduced the number of bystander Tgfbr3^WT^ T_RM_ at the DNFB site, but not local antigen experienced T_RM_ at the VV-OVA site, consistent with our earlier results (**Figure 6I-J**). In contrast, local antigen experienced huNGFR^+^ Tgfbr3^ΔCD8^ T_RM_ from the VV-OVA site that lack TGFβRIII were efficiently depleted from the epidermis by CWHM12 treatment. The depletion of bystander huNGFR^+^ Tgfbr3^ΔCD8^ T_RM_ at the DNFB site was augmented. Notably, numbers of Tgfbr3^ΔCD8^ T_RM_ in cohorts treated with vehicle did not induce a statistically signficant reduction in steady-state T_RM_, indicating that loss of TGFβRIII is not an absolute requirement for T_RM_ epidermal residence in the steady state. Thus, expression of TGFβRIII by local antigen experienced T_RM_ which is downstream of TCR ligation is required for their capacity to remain in the epidermis when activated TGFβ is limiting.

## Discussion

Herein we have demonstrated that the increased capacity of local antigen experienced T_RM_ to persist in the epidermis when levels of TGFβ are limited is mediated by increased expression of TGFβRIII. We also show that local antigen experienced T_RM_ have increased proliferative capacity during repeated antigen recalls. In addition, the increased proliferative capacity was directly correlated with the strength of TCR stimulation during T_RM_ development. Finally, we found that local antigen experienced T_RM_ appear more transcriptionally related to fully differentiated T_RM_. Taken together, these data support a model in which TCR engagement by cognate antigen in the skin is a required final step in T_RM_ differentiation resulting in their increased fitness exemplified by increased proliferative capacity and the ability to persist in the epidermis when active TGFβ is limited.

We propose that the augmented fitness of local antigen experienced T_RM_ represents a mechanism to enrich for high avidity TCR-clones in the epidermis. Skin inflammation recruits T_EFF_ into the skin, some of which develop into T_RM_. In the absence of competition with pre-existing T_RM_, T_RM_ form comparably in the presence or absence of cognate antigen. Thus, we found equivalent numbers of bystander T_RM_ at the DNFB and local antigen experienced T_RM_ at the VV sites with both TCR transgenic and endogenous T cells. In contrast, when new T_EFF_ are recruited into sites with pre-existing T_RM_, there is clonal competition for limited amounts of active TGFβ resulting in enrichment of fitter, local antigen experienced T_RM_ [21]. We now find that this enrichment likely results from 2 different competitive advantages. First, antigen encounter in the skin results in increased expression of TGFβRIII which increases TGFβ avidity for the signaling by the TGFβ receptor. Since TGFβ signaling is required for epidermal persistence, this would provide an advantage for local antigen experienced T_RM_ over bystanders. Second, local antigen experienced T_RM_ have increased proliferation when re-encountering antigen in the epidermis. Following repeated challenges which would be expected outside of SPF conditions, the combination of improved expansion and persistence would work together to enrich for high avidity clones thereby shaping the epidermal CD8^+^ T cell memory pool. Recently, it has been observed that T_RM_ can contribute significantly to the pool of circulating memory cells [55–59]. Thus, mechanisms augmenting epidermal T_RM_ fitness that shape the pool of epidermal T_RM_ may also affect the pool of systemic memory cells and represent an example of extra-thymic clonal section.

When T_RM_ were challenged in a primary antigen recall response, we noted that local antigen experienced T_RM_ expanded to a greater extent than bystanders. This expansion resulted from increased in-situ proliferation with minimal contribution from newly recruited T_EFF_, consistent with prior reports [20, 60, 61]. Interestingly, a similar phenomenon occurred following a second encounter with antigen. This indicates that encounter with peptide at a late time point after T_RM_ differentiation (>50 days) is insufficient to convert bystander T_RM_ into local antigen experienced T_RM_. Thus, there appears to be a window during T_RM_ development when TCR engagement can allow for full differentiation. We also observed after a single recall response that T_RM_ contracted to an elevated baseline suggesting an increase in the epidermal niche. We speculate this may result from a reduced T cell intrinsic requirement for survival and/or homeostatic proliferation factors such as IL-7 or IL-15 or increased expression of these factors by keratinocytes[62, 63]. Altered sensitivity or availability of TGFβ is unlikely to explain the increased niche size as this would be predicted to vary between local antigen experienced and bystander.

Transcriptional analysis of T_RM_ isolated from the small intestine have revealed intra-organ heterogeneity, with unique transcriptional populations arising early during T_RM_ development [64–66]. This aligns well with our identification of 6 distinct transcriptional clusters of epidermal T_RM_. Cluster 3 appears to represent fully differentiated T_RM_ based on comparison with other T_RM_ datasets. In addition, cluster 3 cells more highly expressed the activation and proliferation associated genes *Junb, Fos* and *Dusp1* as well as *Nr4a1*. Increased basal expression of the AP-1 family members *Junb and Fos* could contribute to the enhanced proliferation of antigen-experienced epidermal T_RM_ during a recall response. Intriguingly, memory CD8 T cells lacking the transcription factor Zbtb20 manifest elevated expression of AP-1 family members and mount more robust antitumor responses [71]. The *Nr4a1* gene encodes for Nur77, which is induced by TCR signaling and its expression correlates with peptide avidity. Notably Nur77 is required for T_RM_ formation in the liver [23–25, 67, 68]. Interestingly, cells in cluster 3 only accounted for 27% of T_RM_ that had the opportunity to encounter their cognate antigen in the VV treated flank. We speculate that not all clones at the VV site fully develop into fitter T_RM_ due to lower TCR avidity or specificity to viral antigens only expressed early during infection which would be absent once the clones arrived into skin.

In sum, TCR signaling during T_RM_ differentiation represents a previously unappreciated final step in T_RM_ differentiation. This results in fitter T_RM_ with a lower requirement for TGFβ transactivation due to increased expression of TGFβRIII and enhanced proliferation in response to peptide stimulation. Moreover, the differing responses to altered peptide ligands indicate that the degree of fitness depends on TCR signal-strength. Thus, polyclonal T_RM_ likely develop into a spectrum of bystander to local antigen experienced cells based on TCR avidity. Though we have focused entirely on epidermal T cells, we suspect that these mechanisms may play a role in other epithelial tissues where residency is also dependent upon TGFβ. Additionally, we have solely investigated memory CD8^+^ T cells after acute inflammation, the role of ongoing TCR-engagement during chronic antigen encounter remains unexplored.

## Materials and Methods

### Mice

We generated Thy1.1^+^Rag^−/−^OT-I mice by crossing OT-I mice with Rag^−/−^ and Thy1.1 mice. E8I-*cre*ER^T2^ and ROSA26.LSL.hNGFR reporter mice were developed by Dario A.A. Vignali (University of Pittsburgh) [29]. E8I-*cre*ER^T2^ mice and Thy1.1^+^Rag^-/-^ OT-I mice were crossed with ROSA26.LSL.hNGFR reporter mice to generate *Tgfbr3^WT^* mice and additionally with *Tgfbr3*^loxP^ mice to obtain *Tgfbr3*^ΔCD8^ mice. *Tgfbr3^loxp^* mice were developed by Herbert Y Lin (Program in Membrane Biology/Nephrology, Department of Medicine, Massachusetts General Hospital and Harvard Medical School, Boston, Massachusetts) [69]. We used age- and sex-matched female mice that were between 6 and 12 weeks of age at the start of all experiments. All mice were maintained under specific-pathogen-free conditions and all animal experiments were approved by University of Pittsburgh Institutional Animal Care and Use Committee.

### T_RM_ cell models and blocking TGFβ activation treatments

Mice were infected by skin scarification (skin infection) with 3 × 10^6^ plaque-forming units recombinant vaccinia virus expressing the SIINFEKL peptide of ovalbumin (VV-OVA), or vaccinia virus without recombinant peptide expression (VV). For skin scarification, 45 μL of VV-OVA or VV was applied to shaved left flank (4–5 cm^2^) and the skin were gently scratched 100 times with 27 G needle under anesthesia. VV or VV-OVA infected mice were further treated with 0.15% 1-Fluoro-2,4-dinitrobenzene (DNFB, D1529: Sigma-Aldrich) in 4:1 acetone: olive oil (O1514, Sigma-Aldrich) on the flank opposite the site of infection (40 μL) 5 days post infection. In some experiments, 4:1 acetone and olive oil was added to the stock solution of OVA_257–264_ (SIINFEKL or SIYNFEKL, RP101611, Genscript/Fisher) in DMSO (D2650, Sigma-Aldrich) (10 mg/mL) for a final concentration of OVA_257–264_ at 0.2 mg/mL and 50 μL was painted to the DNFB-treated skin 1 day after DNFB treatment.

### Recall response

50+ days after VV-OVA infection. 4:1 acetone and olive oil was added to the stock solution of OVA_257–264_ (SIINFEKL, RP101611, Genscript/Fisher) in DMSO (D2650, Sigma-Aldrich) (10 mg/mL) for a final concentration of OVA_257–264_ at 0.2 mg/mL and 50 μL was painted to the both flanks.

### CWHM12 treatments

For blocking TGFβ activation with a small molecule integrin inhibitor, compound CWHM12 (kindly provided by Indalo therapeutics, Cambridge, MA) was solubilized in 100% DMSO. Further dilution to 50% DMSO was made in sterile 100% PBS and dosed to 100 mg per kg body weight per day delivered by i.p. injections once a day for 10 days.

### Adoptive transfers

OT-I cells were purified from spleen and lymph nodes by MojoSort Mouse CD8 T Cell Isolation Kit (48007, Biolegend) according to the manufacturer’s instructions and 1 × 10^5^ OT-I cells were intravenously transferred is all experiments. All mice were allowed to rest for 1 day before further experimentation.

### Tamoxifen treatment

Tamoxifen (T5648; Sigma-Aldrich) was dissolved in 1/10th volume of 200 proof ethanol following incubation at 37 °C for 15–30 min with 300 rpm shaking. Corn oil (C8267, Sigma-Aldrich) was added for a final concentration of Tamoxifen at 10 mg/ml and was administered to mice for 5 consecutive days by intraperitoneal injection at 0.05 mg per g body weight, in order to transform approximately 50% of adoptively transferred cells.

### BrdU treatment

BrdU (B5002, Sigma) was dissolved in sterile PBS. 2 mg in 200uL of PBS was injected i.p. daily into each mouse for 2 consecutive days before harvest.

### Immunofluorescence of epidermis

Epidermal sheets were prepared as previously described [30]. Briefly, 2 square cm sections were harvested 2 cm distal from the mid-clavicular line for consistency, and then the epidermal side of shaved defatted flank skin was affixed to slides with double-sided adhesive (3M, St. Paul, MN). Slides were incubated in 10 mM EDTA in PBS for 90 min at 37°C, followed by physical removal of the dermis. The epidermal sheets were fixed in 4% PFA at RT for 15 min. The epidermal sheets were blocked with PBS containing 0.1% tween-20, 2% BSA and 2% rat serum for 1 hour at RT before staining 1 hour with antibodies at RT in PBS containing 0.1% tween-20 and 0.5% BSA. The slides were mounted with ProLong™ Gold Antifade Mount with DNA stains DAPI (P36931, Thermofisher). Images were captured on Keyence BZX800 fluorescent microscope (Keyence, Osaka, Japan). Analysis was performed using BZ-H4A Advanced Analysis Software(Keyence, Osaka, Japan). For enumeration of cells, three images from distant sites randomly determined within an epidermal sheet from a mouse were counted (total 3 mm^2^) manually after image processing by Adobe Photoshop (version 6) and the average number per mm^2^ epidermis was calculated as representative of the epidermal sheet. Anti-CD8α (53–6.7), Thy1.1 (OX-7) and huNGFR (ME20.4), were purchased from Biolegend.

### Depletion of circulation cells

For depletion of circulating memory OT-I cells, OT-I adoptive transfer VV-OVA and DNFB treated mice were injected i.p. with 0.3–1 μg anti-Thy1.1 (HIS51, ThermoFisher) in 200 μL PBS for 10 consecutive days. Depletion of OT-I T cells (< 0.5% of total CD8^+^ T cells and < 1%) were confirmed by staining with anti-Thy1.1 (OX-7) using blood 3 days after depletion. For FTY720 treatment, OT-I adoptive transfer VV-OVA and DNFB treated mice were injected i.p. with FTY720 (10006292, Fisher/Cayan Chemical) at 1 μg/g body weight in sterile PBS with 0.5% DMSO (D2650, Sigma-Aldrich) daily for 10 days. Depletion of circulating cells was confirmed with staining of anti-Thy1.1 (OX-7) and anti-CD8α (53–6.7) using blood 3 days after the end of treatment.

### Flow cytometry

Preparation of single cells suspension from skin, shaved skin was harvested, and fat tissues was removed by forceps mechanically. The skin was then mechanically digested with either scissors or a gentleMACS Dissociator (Miltenyi Biotec, Bergisch Gladbach, Germany), and then resuspended in RPMI 1640 media (Gibco, Grand Island, NY) containing 2.5 mg/ml Collagenase XI (9001-12-1, Sigma-Aldrich), 0.1 mg/ml DNase (04536282001, Sigma-Aldrich), 0.01 M 4-(2-hydroxyethyl)-1-piperazineethanesulfonic acid (7365-45-9, Sigma-Aldrich), and 10% fetal bovine serum, followed by incubation in a shaking incubator for 30 minutes at 37 °C. The resulting cell mesh was filtered through a 40-μm cell strainer (BD Biosciences, San Jose, CA). The single-cell suspension was then resuspended in 44% percoll (89428-524, VWR) in RPMI and layered over 67% percoll in PBS. After 20 minutes of centrifugation at 931 x g at RT, the interface was isolated and resuspended in FACs buffer. LNs (axillary and inguinal) were incubated in 400 U/ml Collagenase D (Roche Applied Science, Penzberg, Germany) and 0.1 mg/ml DNase in RPMI 1640 with 10% fetal bovine serum for 40 minutes at 37 °C and then minced through a 40-μm cell strainer. In some experiments LNs were heat-treated at 60°C in a hot water bath for 1 hour. Blood was collected with heparin (H3393, Sigma-Aldrich) and treated with red blood cell lysing buffer (R7757, Sigma-Aldrich). Single-cell suspensions were blocked with 2.4G2 culture supernatant (ATCC, Manassas, VA). Surface staining was performed in standard FACS buffer for 30 minutes at 4 °C. Anti-CD8α (53–6.7), CD44 (IM7), CD3 (17A2), Thy1.2 (30-H12), Thy1.1 (OX-7), CD69 (H1.2F3), CD103 (2E7), BrdU (3D4), MHC-II (M5/114.15.2), CD45.2 (104) and Annexin V were purchased from Biolegend. Anti-TGFßRIII (1C5H11) was purchased from Sigma-Aldrich. Soluble tetrameric B8R_20–27_/H2-K^b^ complex (made by the NIH Tetramer Core Facility) was conjugated to PE-labeled streptavidin. A BD LSRFORTESSA (BD Biosciences) and Flowjo software (TreeStar, Ashland, OR) were used for analysis.

### Single-cell RNA and Totalseq C-Sequencing - Sample Preparation

#### Flow cytometry and HT antibody staining

Single-cell lymphocyte suspensions were isolated as described above and stained with anti-CD45-APC, anti-CD8a, anti-CD44, anti-CD103 and different TotalSeq-C Anti-Mouse Hashtags (TotalSeq-C Anti-Mouse Hashtags 1-10 Biolegend #155861-155879, in staining buffer (DMEM, 2% FCS and 10mM HEPES)) for 20min at 4°C in the dark. Standard spleen cell suspensions were stained under identical conditions but included anti-CD45-FITC (BioLegend 109806, clone 104) to differentiate them from other samples when pooled. Each sample was labeled with a unique hashtag, enabling the downstream assignment of individual cell (10x cell barcode) to their respective source samples [70].

#### Cell sorting

For each sample, 500-5000K DAPI^-^ CD45^+^ Thy1.2^+^ CD8a^+^ (skin samples) or DAPI^-^ CD45^+^ Thy1.2^+^ CD8a^+^ CD44^+^ CD103^+^ (LN samples) T cells were sorted on the FACSAria III and pooled in a single collection tube at 4C. DAPI was added just before the sort.

#### ImmGenT TotalSeq-C custom mouse panel staining

The pooled single cell suspension was stained with the ImmGenT TotalSeq-C custom mouse panel, containing 128 antibodies (Biolegend Part no. 900004815) and FcBlock (Bio X Cell #BE0307). Because 500,000 cells are required for staining, unstained splenocytes were spiked in to reach a total of 500,000 cells.

#### Second sort

Cells were subsequently sorted a second time with the addition of DAPI to select for live T cells, and include 5,000 total splenocyte standard cells were sorted into a single collection tube.

### Single-cell RNA and TotalseqC-Sequencing - Library Preparation

*Cell encapsulation and cDNA library.* Single-cell RNA sequencing was performed using the 10x Genomics 5’ v2 platform with Feature Barcoding for Cell Surface Protein and Immune Receptor Mapping, adhering to the manufacturer’s guidelines (CG000330). After cell encapsulation with the Chromium Controller, reverse transcription and PCR amplification were performed in the emulsion. From the amplified cDNA library, smaller fragments containing TotalSeq-C-derived cDNA were separated for Feature Barcode library construction, while larger fragments containing transcript-derived cDNA were preserved for TCR and Gene Expression library generation. Library sizes for both cDNA fractions were evaluated using the Agilent Bioanalyzer 2100 High Sensitivity DNA assay and quantified with a Qubit dsDNA HS Assay kit on a Qubit 4.0 Fluorometer.

#### RNA library construction

After enzymatic fragmentation and size selection of the cDNA, the library was ligated to an Illumina R2 sequence and indexed using unique Dual Index TT set A index sequences, SI-TT-B6.

#### TotalSeq-C library construction

Totalseq-C derived cDNA was processed into the Feature Barcode libraries following the manufacturer’s protocol. The library was indexed with a unique dual index TN set TN set A (10x part no. 3000510) index sequence SI-TN-F9.

#### Sequencing

The three libraries were pooled based on molarity in the following proportions: 47.5% RNA, 47.5% Feature Barcode, and 5% TCR. The pooled libraries were sequenced on an Illumina NovaSeq S2 platform (100 cycles) using the 10x Genomics specifications: 26 cycles for Read 1, 10 cycles for Index 1, 10 cycles for Index 2, and 90 cycles for Read 2. Single-cell RNA, TCR and TotalseqC-sequencing - Data processing *Code.* Code is available on https://github.com/immgen/immgen_t_git/

#### Count matrices

Gene and TotalSeq-C antibody (surface protein panel and hashtags) counts were obtained by aligning reads to the mm10 (GRCm38) mouse genome using the M25 (GRCm38.p6) Gencode annotation and the DNA barcodes for the TotalSeq-C panel. Alignment was performed with CellRanger software (v7.1.0, 10x Genomics) using default parameters. Cells were identified and separated from droplets with high RNA and TotalSeq-C counts by determining inflection points on the total count curve, using the barcodeRanks function from the DropletUtils package.

#### Sample demultiplexing

Sample demultiplexing was performed using hashtag counts and the HTODemux function from the Seurat package (Seurat v4.1). Doublets (droplets containing two hashtags) were excluded, and cells were assigned to the hashtag with the highest signal, provided it had at least 10 counts and was more than double the signal of the second most abundant hashtag. Hashtag count data were also visualized using t-SNE to ensure clear separation of clusters corresponding to each hashtag. All single cells from the gene count matrix were uniquely matched to a single hashtag, thereby linking them unambiguously to their original sample.

#### Quality control (QC) and batch correction

Cells meeting any of the following criteria were excluded from the analysis: fewer than 500 RNA counts, dead cells with over 10% of counts mapping to mitochondrial genes, fewer than 500 TotalSeq-C counts, or positivity for two isotype controls (indicating non-specific TotalSeq-C antibody binding). Non-T cells were excluded based on the expression of the MNP gene signature, B cell signature, ILC gene signature, and the absence of T cell gene signature (score calculated using AddModuleScore_UCell). CITEseq data did not meet quality control and was not used in the analysis.

#### ImmGen T integration

The data was integrated with the rest of the ImmgenT dataset using the SCVI.TOTALVI model (v1.2.0) and the 10x lane as a batch parameter. Dimensionality reduction was performed using the pymde.preserve_neighbors() function with default parameters [CITATION: https://pymde.org/citing/index.html]. Cell clustering was carried out using the FindClusters() function in Seurat. Manual annotation by the immgenT consortium was done using protein and RNA expression of Cd3e, Trbc1, Trbc2, Cd4, Cd8a, Cd8b1, Foxp3, Mki67, Sell, Cd44, Trgc1, Itgax, Itgam, Ms4a1, CD3, TCRB, THY1.2, CD4, CD8A, CD8B, CD62L, CD44, TCRGD, TCRVG1.1, TCRVG2, TCRVG3, CD19, CD20, ITAM.CD11B, ITAX.CD11C, KLRBC-NK1.1. Data was visualized using the Rosetta software (https://cbdm.connect.hms.harvard.edu/ImmgenT/PublicRosetta/). ImmgenT integration and annotation available on https://www.immgen.org/ImmGenT/.

#### Clustering and dimensionality reduction

Using Seurat v4.2. [https://pubmed.ncbi.nlm.nih.gov/31178118/], the variance-stabilizing transformation (VST method) was applied, and PCA was conducted on the top 2000 genes. Principal components explaining 80% of the total variance were selected for two-dimensional reduction using UMAP. Clustering was performed on these principal components using the FindClusters() function in Seurat.

#### Differential Expression

To determine differentially expressed genes, FindMarkers() within Seurat was used. AddModuleScore() was used to visualize aggregated expression of a set of genes. Volcano plots were created using EnhancedVolcano (v1.14.0).

### Signature score analysis

Signature score was calculated by first generating differentially expressed genes of each cluster or condition compared to naïve T cells from the spleen control, and then calculating an average z-score for every up and down gene in a core gene signature datasets. The core gene datasets were generated by Jaiswal et al. [35] by calculating the exclusive expression of genes from T-cells in the datasets GSE47045 [23], GSE10239 [37] and GSE41867 [38]. Comparative programs for T_RM_ development over time were analyzed against the transcripts taken from the top 100 differentially expressed genes of each T_RM_ timepoint/ naïve from GSE79805 [39] by adapting published scoring methods with this data set Jaiswal et al. [35]. The overall signature score was scored based on expression data that was quantified using RSEM in transcripts per million (TPM), then log-transformed as log2(TPM+1). It was then centered for each gene across all cells. The centered data was averaged across sets of genes to define signature scores. We subtracted a control score from the signature score, which is defined using the same process on randomly selected gene-sets. A randomized control was generated by the average enrichment of each group compared to a randomly generated gene set of an equal number of probes, as iterated Jaiswal et al. [35].

### Quantification and statistical analysis

Groups were compared with Prism software (GraphPad) using two-tailed paired Student’s t-test for comparison of left and right flanks, two-tailed unpaired Student’s t-test for comparison between animals, or Dunnett’s test for comparisons of more than two groups. Data is presented as each data point and mean with the standard error of the mean (s.e.m.). p < 0.05 was considered significant.

## Acknowledgements

We thank Daniel Bernard (Division of Endocrinology and Metabolism, McGill University, Montreal, Canada) for providing the Tgfbr3^fl/fl^ mice. We thank the members of the Kaplan and Vignali laboratories and members throughout the departments of Dermatology and Immunology for helpful discussions. We also thank the Division of Laboratory Animal Resources of the University of Pittsburgh for excellent animal care. The experimental designs were created with BioRender.com.

## Author contributions

Eric S. Weiss, Toshiro Hirai and Daniel H Kaplan designed and interpreted experiments. Eric S. Weiss, Toshiro Hirai, Haiyue Li, Andrew Liu, Shannon Baker, Ian Magill, Jacob Gillis, Youran R. Zhang, Torben Ramcke and Kazuo Kurihara performed experiments. Eric S. Weiss and David Zemmour conducted bioinformatics analysis. David Masopust, Niroshana Anandasabapathy, Harinder Singh, David Zemmour and Laura Mackay provided technical and conceptual assistance. Eric S. Weiss and Daniel H. Kaplan wrote the manuscript, and all authors edited it.

