## Supplementals figures 1 and 2 for "Epidermal Resident Memory T Cell Fitness Requires Antigen Encounter in the Skin"

**Supplemental figure 1:** (A) Quantification of percent $CD103^+$ T cells from VV and DNFB treated flanks, gated on live $CD45.2^+CD8^+CD90.2^+$. Each bar represents the mean +/- SEM for n= 6 animals. (B) Gating strategy for skin single-cell RNA-seq. (C) Flow plots of CD69 and CD103 expression of VV and DNFB treated flanks for single-cell RNA-seq gated as in (B). (D) Heatmap showing the top 10 DEG per cluster when compared with each other, calculated by FindAllMarkers of Seurat V2, percent expressed > 0.02, Log2foldchange > 2.

Weiss et al. Supplemental Figure 1

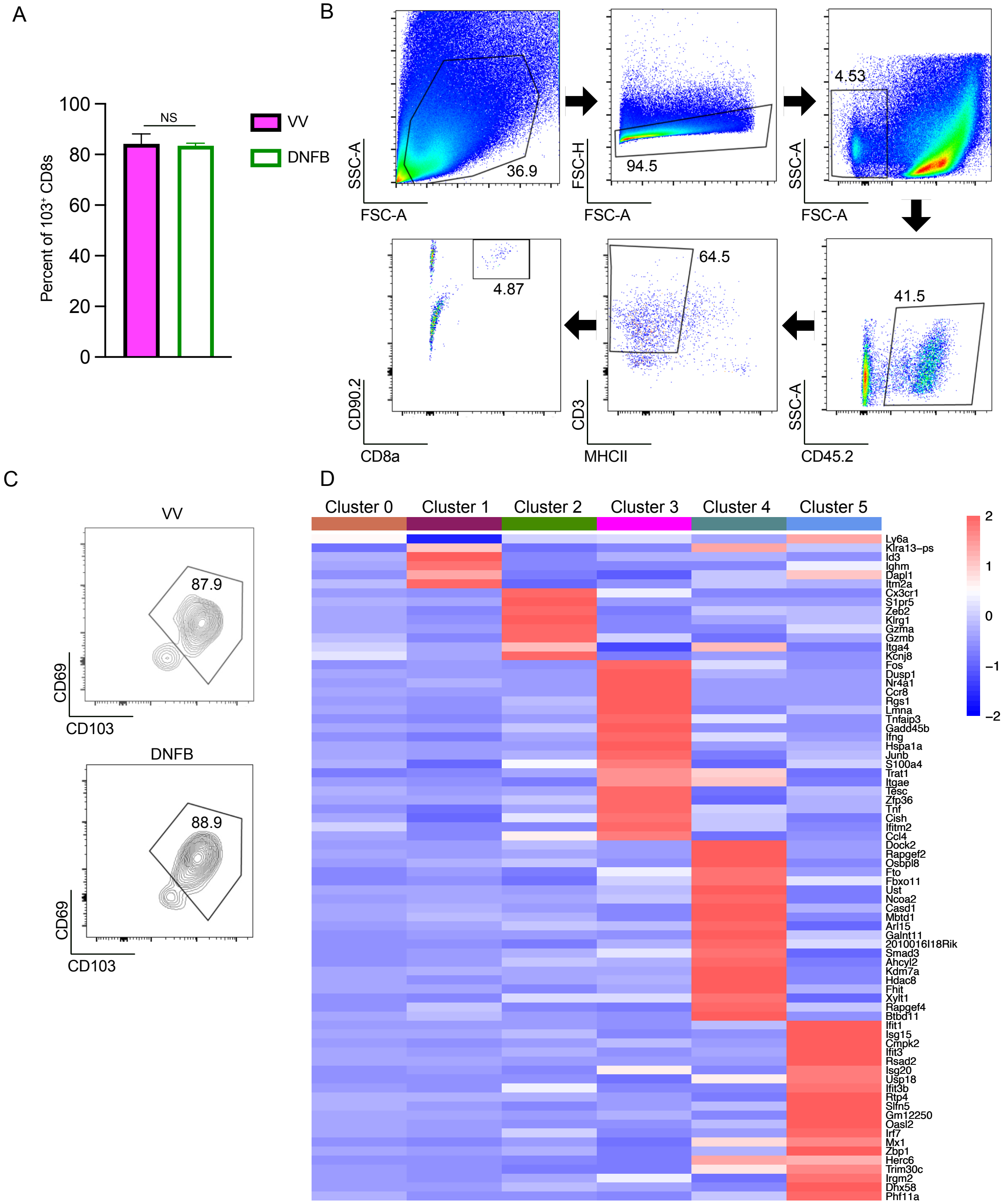

**Supplemental figure 2:** (A) Representative flow plots of blood from Thy1.1 OT-I adoptive transferred mice after 6 days of vehicle or FTY720 treatment, gated on liveCD45$^+$. (B) Quantification of total Thy1.1 OT-I in blood after FTY720 treatment, gated as in (A). Each bar represents the mean +/I SEM for n= 6 animals. (C) Representative flow plots of blood of CD45$^+$CD3$^+$CD8$^+$ cells after isotype or Thy1.1 depleting antibody treatment. (D) Quantification of Thy1.1 OT-I in blood before and after isotype or anti-Thy1.1 depleting antibody treatment, gated as in (C). (E) Representative epidermal whole mount of Thy1.1$^+$ cells in the epidermis at steady state of VV-OVA or DNFB treated flanks after 6 days of isotype or anti-Thy1.1 depleting antibody treatment. Each symbol represents data from an individual animal. Data is representative of 3 separate experiments. *$p<0.05$ by unpaired Student's t-tests. Scale bar represents 50um.

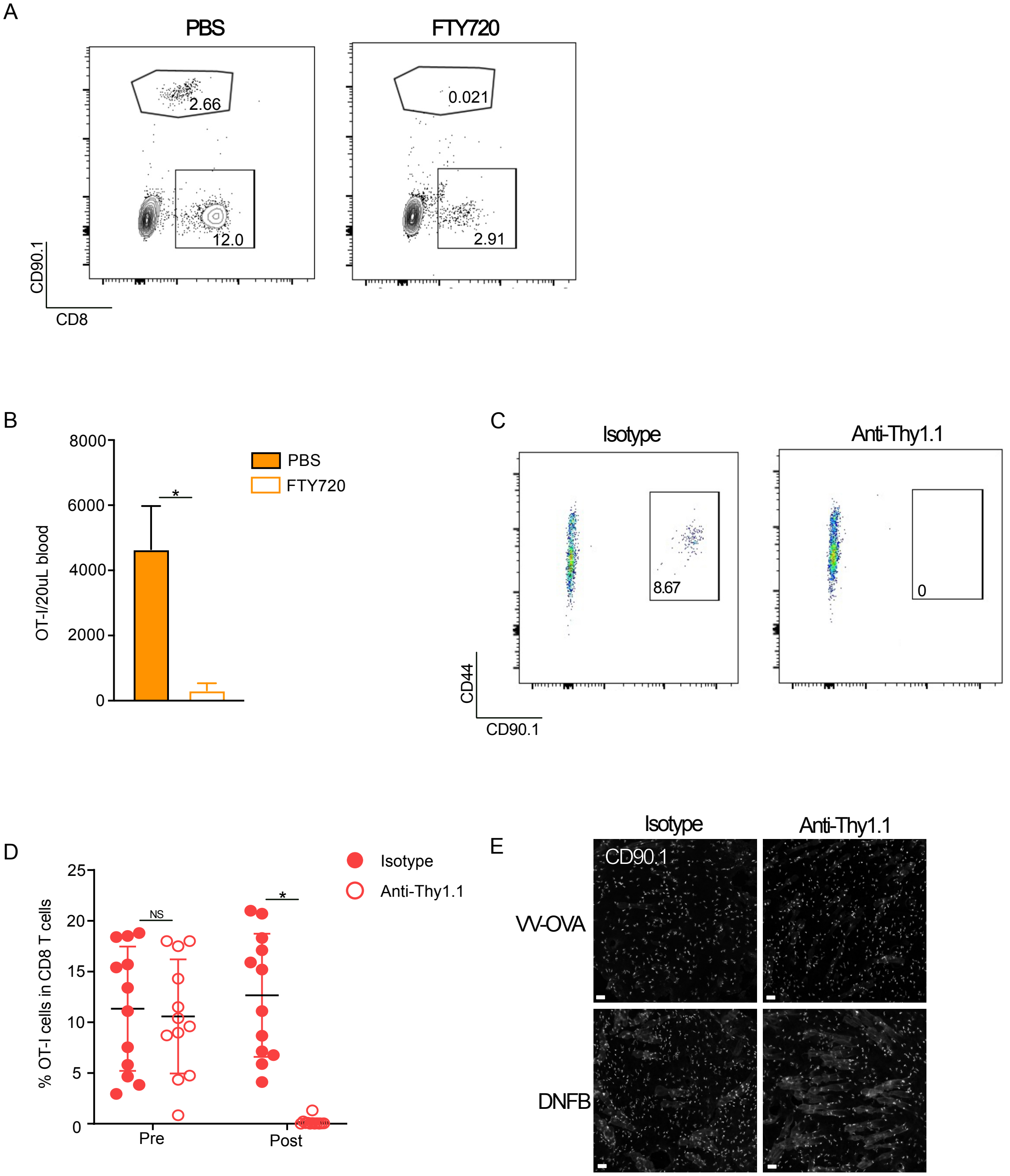
A
PBS
FTY720
2.66
12.0
0.021
2.91
CD90.1
CD8
B
8000
6000
4000
2000
0
OT-I/20uL blood
*
PBS
FTY720
C
Isotype
Anti-Thy1.1
8.67
0
CD44
CD90.1
D
25
20
15
10
5
0
% OT-I cells in CD8 T cells
NS
*
Pre
Post
Isotype
Anti-Thy1.1
E
Isotype
Anti-Thy1.1
CD90.1
VV-OVA
DNFB
